# Impact of the rotational speed and counter electrode configuration on the performance of a rotating disc bioelectrochemical reactor (RDBER) operated as microbial electrolysis cell

**DOI:** 10.1101/2025.03.06.641858

**Authors:** Zhizhao Xiao, Max Rümenapf, Max Hackbarth, Andrea Hille-Reichel, Harald Horn, Johannes Eberhard Reiner

## Abstract

A 10 L Rotating Disc Bioelectrochemical Reactor (RDBER) was operated as a microbial electrolysis cell (MEC) under different rotational speeds and counter electrode configurations. Increasing the anode’s speed from 0.25 to 1 rpm raised the anodic current density from 55 ± 14 to 100 ± 7 A m^-3^ while increasing hydrogen production rates from 0.05 ± 0.01 to 0.18 ± 0.01 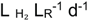. Higher speeds provided no further benefit. Moving the counter electrodes to the upper reactor half reduced observed hydrogen shuttling. The modified RDBER reached current densities of 1.98 ± 0.11 A m^-2^ and 0.99 ± 0.03 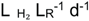 hydrogen production. Optical coherence tomography confirmed biofilm morphology changes but no significant increase in biovolume or substratum coverage. Hydrogen recovery remained below 50%. While the RDBER achieved high volumetric current densities and volumetric hydrogen production rates compared to other MEC pilots, improvements in anodic current density and cathodic hydrogen recovery are required for practical application.

**Graphical Abstract:** 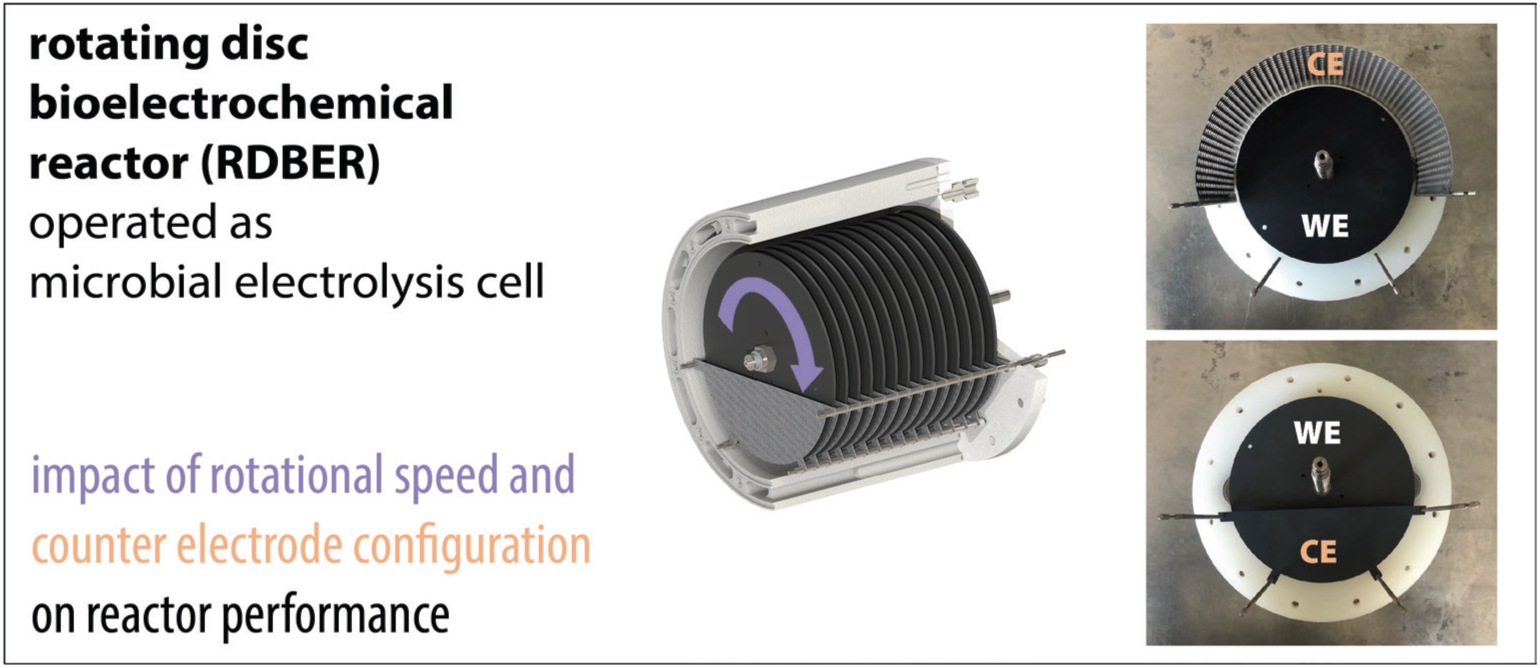

**Highlights:**

- 10 L RDBER was operated as MEC in batch experiments at different rotational speeds
- hydrogen shuttling was reduced through cathode displacement
- Current densities of 198 ± 11 A m^-3^ and H_2_-production rates of 0.99 ± 0.03 L L_R_^-1^ d^-1^
- Anodic biofilm parameters were not significantly altered by cathode modification

## 1. Introduction

Microbial electrochemical technologies (METs) intertwine applied electrochemistry with microbial metabolism, either directly or indirectly (Schröder et al., 2015). METs are receiving research attention mainly due to their potential contribution to electricity-driven waste stream transformations within a future sustainable economy (Ieropoulos et al., 2024; Jiang et al., 2023; Jourdin and Burdyny, 2021). A specific implementation of this technology is a microbial electrolysis cell (MEC). Here, the exergonic microbial degradation of organic compounds is electrically linked to an abiotic, cathodic hydrogen production (Rozendal et al., 2006). The electrical energy input required to drive the cathodic hydrogen evolution reaction (HER) can be significantly lower in a MEC compared to conventional water electrolysis (Kim et al., 2025). In this way, MECs have positioned themselves at the forefront of the evolving field of METs - by coupling the treatment of carbon-rich waste streams with the surplus of an electrocatalytic hydrogen production. Microbial electrochemical technologies have undergone extensive laboratory study over the last decades and are currently finding their way into pilot-scale implementations (Das and Ghangrekar, 2024; Ieropoulos et al., 2024). Inevitably, this also drives the development of novel reactor designs to meet the scaled demands of a pilot plant (Enzmann et al., 2019). At the laboratory scale (<1 L reactor volume), zero-gap flow cells clearly outperform other reactor concepts in the pursuit of increased current densities and enhanced hydrogen production rates (Kim et al., 2025; Rossi et al., 2023, 2022). These zero-gap designs are optimized versions of a conventional dual-chamber MEC reactor, with a membrane separating anode and cathode chambers to ensure hydrogen purity and prevent electrochemical short circuits (Gautam et al., 2023; Liu et al., 2010). However, while the ongoing development of zero-gap flow cells partly addressed challenges associated with conventional membrane-based MEC designs – such as the formation of pH gradients and increased internal resistances – high operational costs related to maintenance and membrane fouling might remain a significant issue (Bian et al., 2024; Kim et al., 2025; Kracke et al., 2018; Utesch and Zeng, 2018). Noteworthy, the outstanding performance of these systems is currently limited to small-scale reactors (a few milliliters in anodic working volume), which appear to be at least challenging to scale linearly. Hence, it is often proposed that the increase in reactor working volume and electrode surface area should be achieved through numbering up of small zero-gap reactors (Enzmann et al., 2019; Jiang et al., 2023). The modular design of commercial-scale water electrolyzers is often held up as a reference, as a similar stacking concept is already practiced here. Yet, connecting thousands of individual modules in order to provide sufficient anode working volume required to treat real liquid waste streams may result in a plethora of potential leakage points which considerably complicates periodic inspection and maintenance during long-term operation. Besides, prolonged narrow flow channels could also lead to high flow resistances leading to pressure drops that might result in uneconomical pumping energy demands. Additionally, the narrow flow channels of current zero gap designs may be prone to clogging due to biofilm formation and/or suspended solids in real waste streams (Kim et al., 2025). Finally, the distribution of inter-cell voltages in large stacks remains unknown, as do possible pH and nutrient gradients that may develop within or between individual stacks.

Taken together, while clearly demonstrating the most promising performance on a small scale, numbered-up dual-chamber MECs may not yet be suitable for all large-scale MET applications, such as treating high volumetric waste streams with an elevated content of suspended solids. Hence, novel creative reactor concepts that circumvent these limitations of conventional dual chamber MEC-architectures could facilitate a broader implementation of METs beyond the laboratory, enabling practical deployment in large-scale waste treatment and bioelectrochemical production.

Along these lines, a novel reactor design - the Rotating Disc Bioelectrochemical Reactor (RDBER) - has recently been introduced (Hackbarth et al., 2023). The RDBER employs 14 rotating graphite discs as working electrodes, providing a total working electrode surface area of 1 m2 within a 10 L reactor volume, thereby offering a convenient ratio of working electrode surface area to reactor volume of 100:1. The RDBER design allows for easy scaling while maintaining the surface-to-volume ratio for example by elongating the reactor chamber, thereby increasing the number of graphite discs that can be mounted on the rotating shaft (Pott, 2023). The reactor is fully autoclavable, can be chemically sterilized in situ and is thus, in combination with the integrated temperature control, especially suitable for axenic cultivations. As a single-chamber BES, the RDBER deliberately avoids the use of membranes. While simplifying the reactor design and avoiding complications caused by membrane fouling and membrane failure on the one hand, such a single-chamber system can be prone to electrochemical short circuits, that can result in decreased efficiencies compared to a conventional two-chamber set up.

The RDBER builds upon the principles of so-called Rotating Biological Contactors (RBCs), which are widely used in wastewater treatment (Hassard et al., 2015). RBCs consist of closely spaced discs that serve as biofilm substratum, mounted on a rotatable horizontal shaft. These reactors are characterized by low maintenance and operating costs, ease of process control, and a compact design (Cortez et al., 2008). The rotation of the biofilm-covered discs through the bulk medium and the gas phase of the reactor ensures the supply of substrate and nutrients to the biofilm, thereby reducing the reliance on energy-intensive pumping or stirring. While a faster rotation counteracts the limiting diffusive transfer of nutrients and substrate into the biofilm, it also directly increases the shear force acting on the biofilm and influences the overall energy consumption of the system. Therefore, the rotational speed of the working electrodes can be defined as a key operating parameter of the RDBER.

Although the RDBER was originally designed as a platform for a thermophilic, oxic microbial electrosynthesis process, both anodic and cathodic biofilms have been successfully cultivated in the reactor in proof-of-concept studies (Hackbarth et al., 2023; Juergensen et al., 2025; Knoll et al., 2023). However, systematic research evaluating the potential and limitations of the reactor as a MEC pilot is lacking - a fundamental prerequisite for further optimization of the design. In order to evaluate its performance and assess whether the RDBER can compete with other MEC upscale attempts, the model exoelectrogens *Shewanella oneidensis* and *Geobacter sulfurreducens* were cultivated in co-culture at different working electrode speeds in a series of batch experiments. Since considerable hydrogen shuttling from the cathode into the anodic biofilm was observed, we consequently attempted to mitigate this electrochemical short circuit by modifying the design of the cathode configuration in the RDBER. Finally, we compare the key performance indicators of the optimized reactor with those of other MEC scale-up attempts in the range of 10 L and above.

## 2. Material and methods

### 2.1 Strains and preculture conditions

*Shewanella oneidensis* MR-1 was routinely cultured aerobically in LB-Lennox-Medium (at 30 °C, 180 rpm), while *Geobacter sulfurreducens* PCA was pre-grown under anoxic conditions at 30 °C in BES-Medium (pH 7.4). The detailed composition of this minimal medium can be found elsewhere (Hackbarth et al., 2023). Briefly, 20 mM sodium acetate and 40 mM of disodium fumarate were added as electron donor and acceptor, respectively. Mid-log phase cells were harvested and washed twice in BES-Medium omitting fumarate before final inoculation of the reactor.

### 2.2 Reactor design

A detailed description of the design and configuration of the bioelectrochemical system employed for all experiments in this study has been provided in a previous publication (Hackbarth et al., 2023). The reaction chamber of this rotating disc bioelectrochemical reactor (RDBER) has an inner volume of 10 L. Relative to the working electrode area of 1 m^2^ this results in a surface-to-volume ratio of 100 m^2^ m^-3^. The working electrode consists of 14 graphite discs - each with a diameter of 210 mm - mounted on a rotatable titanium shaft, which also serves as current collector. A high-resolution stepper motor on the back of the reactor (Lexium MDrive Motion Control, Schneider Electric) allows precise, load-independent control of the rotational speed of the titanium shaft. In its original configuration, the counter electrode consisted of 15 semicircular mixed metal oxide (MMO) coated titanium plates (platinode type 177, Umicore) that were mounted on four threaded titanium rods located in the lower half of the cylindrical reaction compartment (Fig. 1a shows a detailed view of a single assembled titanium plate on top of a single graphite disc anode). The individual half-disc-shaped counter electrodes were each mounted alternately between the working electrode discs with a distance of 5.5 mm to the two adjacent discs. The active counter electrode area added up to approximately 0.5 m^2^.

**Fig. 1:**
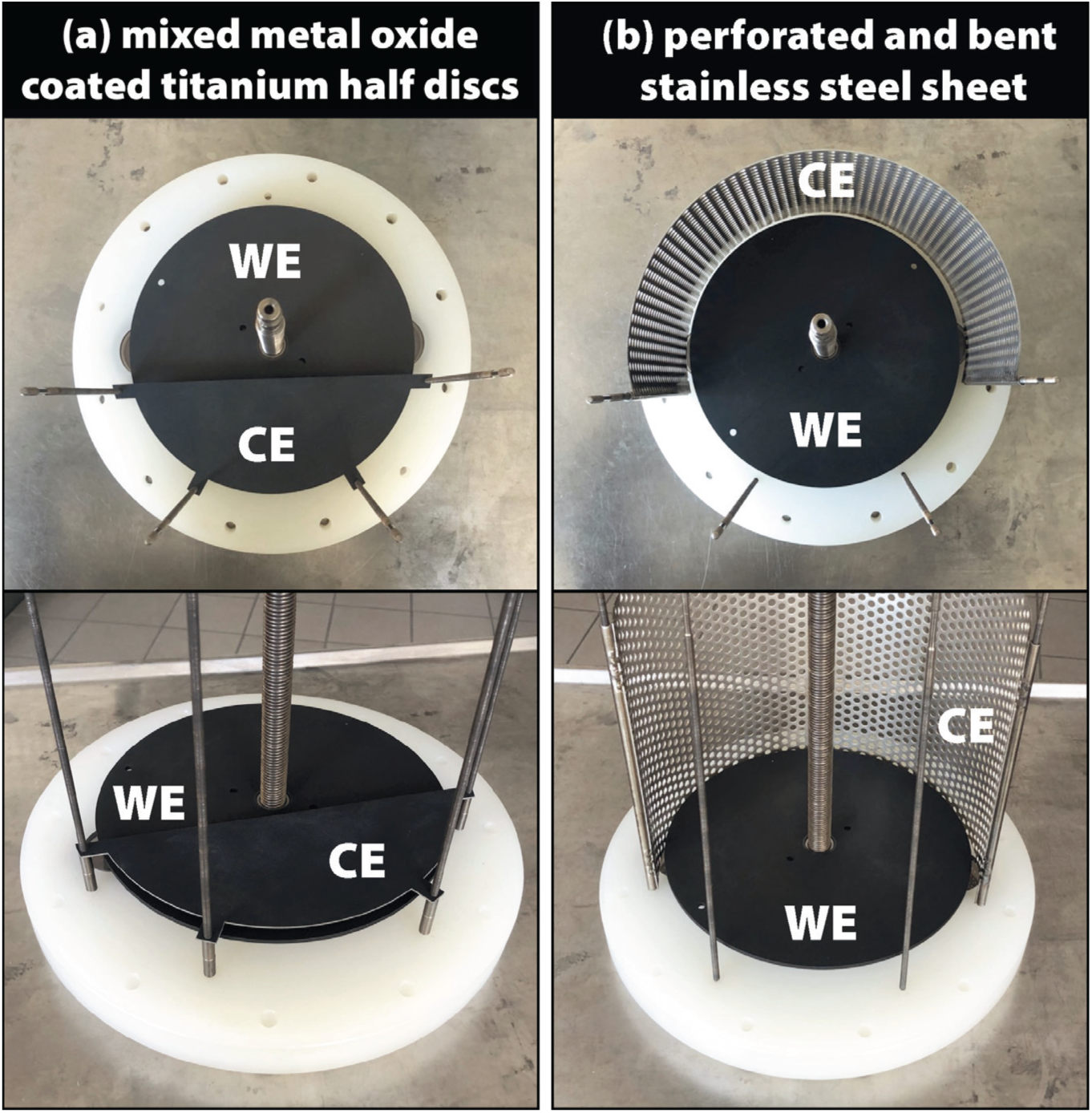
Photographs of the different counter electrode (here the cathode) configurations of the RDBER used in this study. For clarity, only one of the 14 working electrode discs was installed on each picture. (a) semicircular MMO-coated titanium counter electrodes located in the lower half of the reactor (exemplary the photo depicts only one of the 15 counter electrodes). (b) bent and perforated stainless-steel sheet (grade 1.4571) serves as optimized counter electrode placed above the rotating working electrodes. WE – Working electrode; CE – Counter electrode.

To minimize observed hydrogen-shuttling in between the original titanium counter electrodes and the biofilm on the working electrode, a new counter electrode made from a bent stainless-steel (1.4571) sheet was examined in this study (Fig. 1b). For this purpose, a pair of stainless-steel pipes were first screwed onto the two upper titanium rods that originally served for the mounting of the original titanium electrodes. The stainless-steel sheet was bent into a round shape and welded to the stainless-steel pipes so that it was now located on top of the anodes in the upper half of the reaction chamber. A perforation in the electrode sheet allows emerging hydrogen to easily escape towards the gas outlet in the upper part of the reactor and is intended to prevent evolving hydrogen bubbles from accumulating under the steel sheet. The active surface area of the new stainless-steel counter electrode amounted to 0.111 m^2^.

### 2.3 Reactor operation

Prior to each new start-up of the reactor, the system was completely disassembled into its individual parts in order to clean them successively with isopropanol, ethanol and demineralized water. Subsequently, the reactor was reassembled and sterilized by autoclaving. The reactor was then filled with approximately 10 L of previously inoculated BES medium (pH 7.4, 20 mM sodium lactate and 20 mM sodium acetate as electron donors). This involved inoculation with *S. oneidensis* and *G. sulfurreducens* in a 9:1 ratio, resulting in an initial OD_600_ of 0.1. The medium was recirculated at a pump rate of 500 mL min^-1^. A heating jacket in the reactor shell enabled a circulation thermostat-dependent control of the reactor temperature, which was kept constant at 30 °C. By means of a potentiostat (Interface 5000P, Gamry Instruments, Warminster, USA), a constant potential of 0 mV vs. SHE (standard hydrogen electrode) was maintained at the working electrode for the duration of a single experiment (9 days each) while recording the current densities measured throughout an experiment. The evolving gases (e.g. H_2_ and CO_2_) were collected in a gas sampling bag after passing a MilliGascounter (Ritter, Germany) for measurement of the volumetric flow rate. Analysis of the collected gases was conducted by means of a Micro GC (490 Micro GC, Agilent Technologies, Germany) loaded with two carrier gases (argon and helium) and two stationary phases (CP-Molsieve 5 Å Plot and Por-aPLOT U J&W GC columns, Agilent Technologies, Germany). Finally, media samples were taken twice a day in order to determine the concentration of organic acids by means of a Metrohm 881 Compact Pro Ion Exchange Chromatograph with a Metrosep Organic Acids 250/7.8 column (Metrohm, Switzerland). At the end of each experiment after 9 days, optical coherence tomography (OCT) was used for 3D visualization of the anodic biofilm, using a GANYMEDE II spectral-domain OCT system (Thorlabs GmbH, Dachau, Germany) with an LSM04 objective. The digital image processing of the OCT data sets and the calculation of the biofilm structure parameters mean biovolume 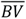 (µm^3^ µm^-2^) and mean substratum coverage 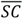 (−) were conducted using imageJ (Bauer et al., 2019; Schneider et al., 2012) and in-house made macros (Bauer et al., 2019) as described elsewhere (Hackbarth et al., 2023). All experiments were performed at least in duplicate.

### 2.4 parameters for reactor performance characterization

The Coulombic efficiency *CE* (%) of an anodic half-cell reaction was calculated as the ratio of the moles of electrons transferred to the anode calculated from the working electrode current densities *n*_*e*(*cd*)_, and the moles of electrons *n*_*e*(*oa*)_ theoretically released from the full oxidation of the consumed organic acids – where the consumption is based on the collected ion chromatography data:

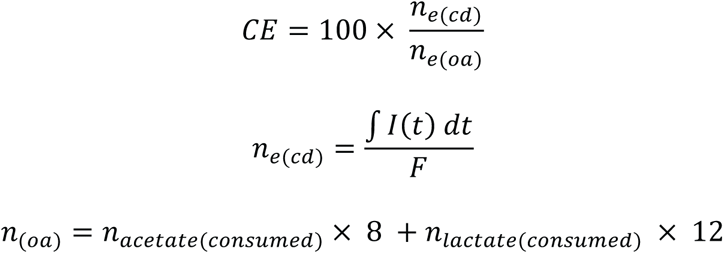

where *F* is the Faraday constant and *I* (A) describes the measured anodic current.

The hydrogen yields 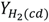 – in literature often referred to as cathodic hydrogen recovery or as r_cat_ – and 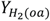 (%) were calculated as the ratio of the actually produced moles of hydrogen measured at the gas outlet of the reactor 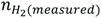, and the moles of hydrogen that theoretically could have been produced based on the measured current densities as described in 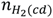 or from the electrons theoretically released during a full oxidation of the degraded organic acids, defined as 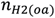.

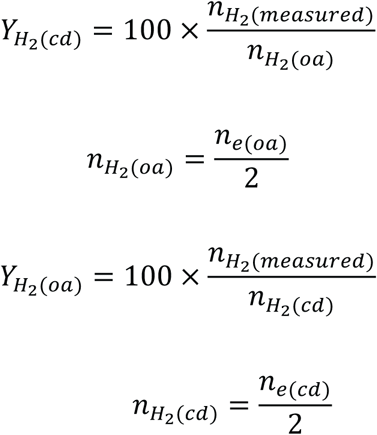

## 3. Results and discussion

### 3.1 Impact of the rotational speed on reactors performance

As part of the iterative process of exploiting the RDBERs potential, the impact of the anodes’ rotational speed on the performance of the reactor operated as a microbial electrolysis cell was assessed in a series of batch experiments. In this context, the RDBER was operated at constant anodic rotational speeds (0.25, 1 and 2 rpm) for a period of 9 days each, while recording key parameters such as current densities, hydrogen production rates, and organic acid degradation. An axenic co-culture of *Geobacter sulfurreducens* and *Shewanella oneidensis* served as inoculum in all experiments. The working electrode potential was kept constant at 0 mV vs. SHE by means of a potentiostat. As a result, the slowest rotational speed tested (0.25 rpm) led to anodic current densities of up to 0.55 ± 0.14 A m^-2^ (corresponds to a volumetric current density of 55 ± 14 A m^-3^). As depicted in Fig. 2, by increasing the rotational speed to 1 rpm, maximum current densities of up to 1 ± 0.07 A m^-2^ (100 ± 7 A m^-3^) could be achieved.

**Figure 2:**
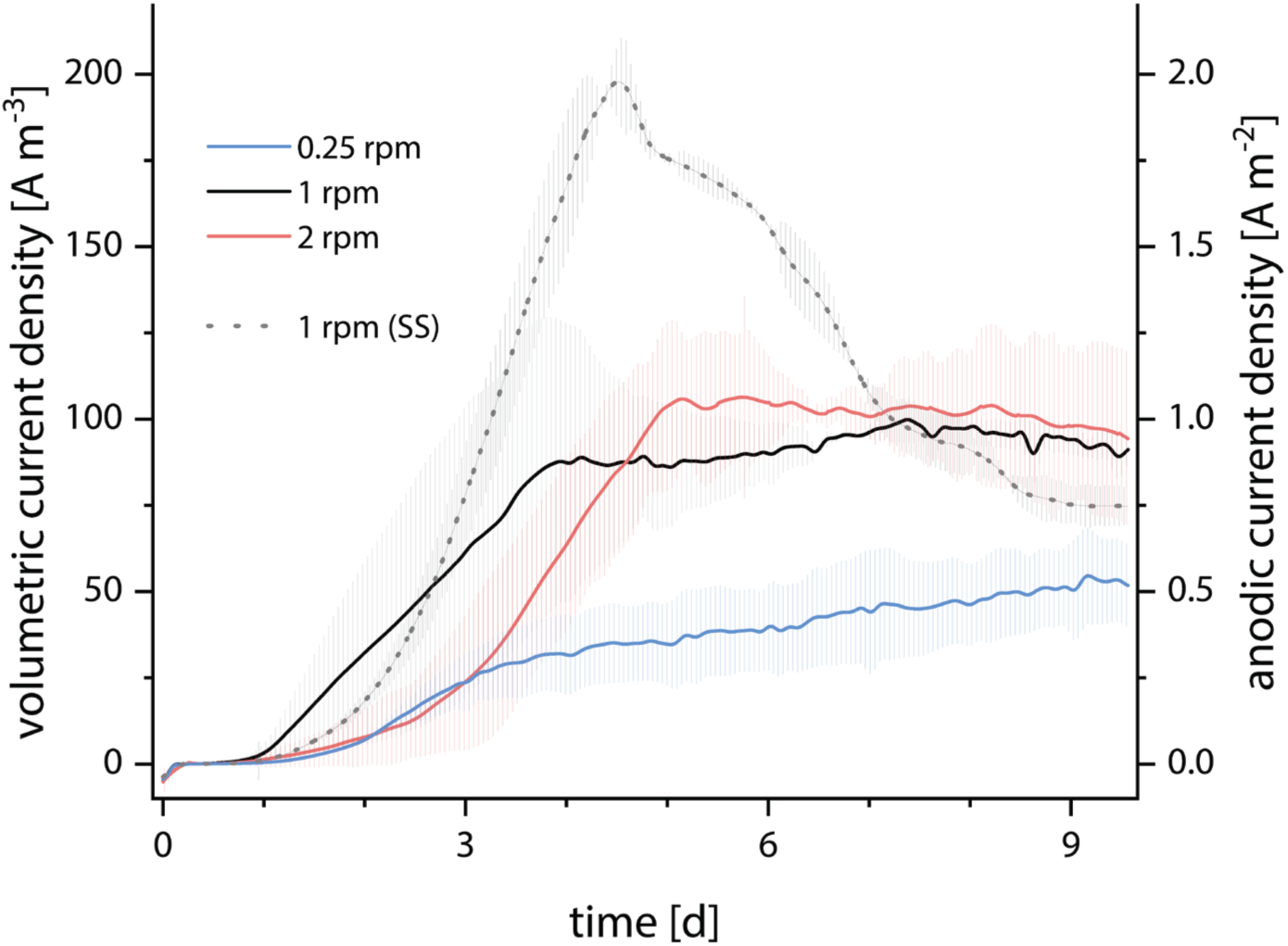
Impact of the rotational speed of the working electrodes on the volumetric and anodic current densities over time (straight lines). The dotted line corresponds to the measured current densities of the improved reactor system with the optimized stainless-steel (SS) counter electrode at 1 rpm.

A further doubling of the speed to 2 rpm did not significantly increase the obtained current densities. Rather, a prolonged initial ‘lag phase’ was observed in the current density profile of the 2 rpm experiments. The degradation profile of the respective organic acids (see Fig. 3a and 3b) showed a similar correlation between rotational speed and performance. By increasing the rotational speed from 0.25 rpm to 1 rpm, acetate degradation rates approximately tripled (from 14.9 ± 3.6 to 45.3 ± 2.2 mmol d^-1^ m^-2^ for the maximum and from 7.3 ± 1.7 to 22.3 ± 0.3 mmol d^-1^ m^-2^ for the average degradation rate). Accompanying this, the maximum and average lactate consumption rates by *S. oneidensis* more than doubled up to 19.9 ± 4.3 mmol d^-1^ m^-2^ (maximum) and 8.1 ± 1.0 mmol d^-1^ m^-2^ (average) respectively. Operating the RDBER at 2 rpm did not result in any further significant difference in the organic acids degradation rates compared to the experiments conducted at 1 rpm. Another process parameter that was heavily affected by the speed of the working electrodes was the hydrogen production rate. Again, the most notable difference in both the average and maximum hydrogen production rates was evident between the 0.25 rpm and 1 rpm experiments (See Fig. 3c and 3d). Here, the maximum hydrogen production rate was increased by a factor of four to slightly exceed 0.4 liters per liter of reactor volume and day. Likewise, the average daily hydrogen production quadrupled from 0.05 ± 0.01 to 0.18 ± 0.01 liters per liter of reactor volume. Increasing the anodes speed to 2 rpm did not significantly change the average or maximum hydrogen production rates. Rittmann et al. employed a rotating biological contactor (non-electrified) to assess oxygen mass transfer into an aerobic biofilm growing on non-conductive discs (Rittmann et al., 1983). However, it is important to note that the Rittmann-RBC was operated as a gas-liquid contactor, exposing half of the disc surface to air, which may limit the comparability of their results with our study. Nevertheless, they demonstrated an increase in aeration with increasing rotational speeds up to 30 rpm. In our study, since acetate was likely not limiting, one of the primary effects of rotation during MEC operation could have been enhanced mixing and the faster removal of anodically produced protons. This effect improved as the rotational speed increased from 0.25 to 1 rpm, but no further improvement was observed beyond that speed. It should also be mentioned that no abiotic organic acid degradation or hydrogen production was measured in a sterile control experiment, nor could any influence of the rotational speed on the anodic idle current of the non-inoculated reactor be observed (*see Supplementary Figure 1*). Based on the production rates described above, the respective yields of the RDBER operated as a microbial electrolysis cell were determined. Coulombic “efficiencies” (*CE*) were calculated as the ratio of electrons transferred to the anode (based on anodic current densities) and electrons released through the oxidation of the degraded organic acids (measured by means of ion chromatography). The *CE*s obtained in this manner (see Fig. 3e) ranged well above 100 % (up to 353 ± 3 % in the 0.25 rpm run). However, to be consistent with the law of conservation of energy, “efficiencies” above 100 % should be critically scrutinized and can be attributed to the fact that not all electron sources available to the anodic biofilm were included in the above calculation. Since care was taken to ensure that the employed minimal medium did not contain any respiratory electron donors accessible to the co-culture, the hypothesis was made that some cathodically produced hydrogen was re-oxidized by the anodic biofilm, resulting in an additional source of anodic electrons. Analyzing the calculated hydrogen yield 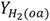 seems to support this assumption of hydrogen shuttling between the two electrodes: Only around 60 % of the hydrogen, theoretically expected to be cathodically produced as a result of the complete anodic oxidation of the consumed organic acids (not accounting for biomass as a carbon and electron sink), was detected at the gas outlet of the reactor - with a negligible influence of the rotational speed (Fig. 3e). Although this hydrogen loss could also be caused by leaks in the reactor itself, hydrogen yield 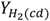, calculated based on the moles of hydrogen that theoretically should have been produced based on the measured current densities, amounted to only around 20 %, which suggests that some of the hydrogen is re-oxidized by *G. sulfurreducens* in the anodic biofilm and thus remains trapped in a ‘hydrogen loop’ in the reactor. This short-circuiting of a microbial electrolytic cell, which results in a significant reduction in rates and yields of the desired electrochemical process, has previously been observed in other studies as a typical unintended side effect of a membrane-less bioelectrochemical system (Wang et al., 2023).

**Figure 3:**
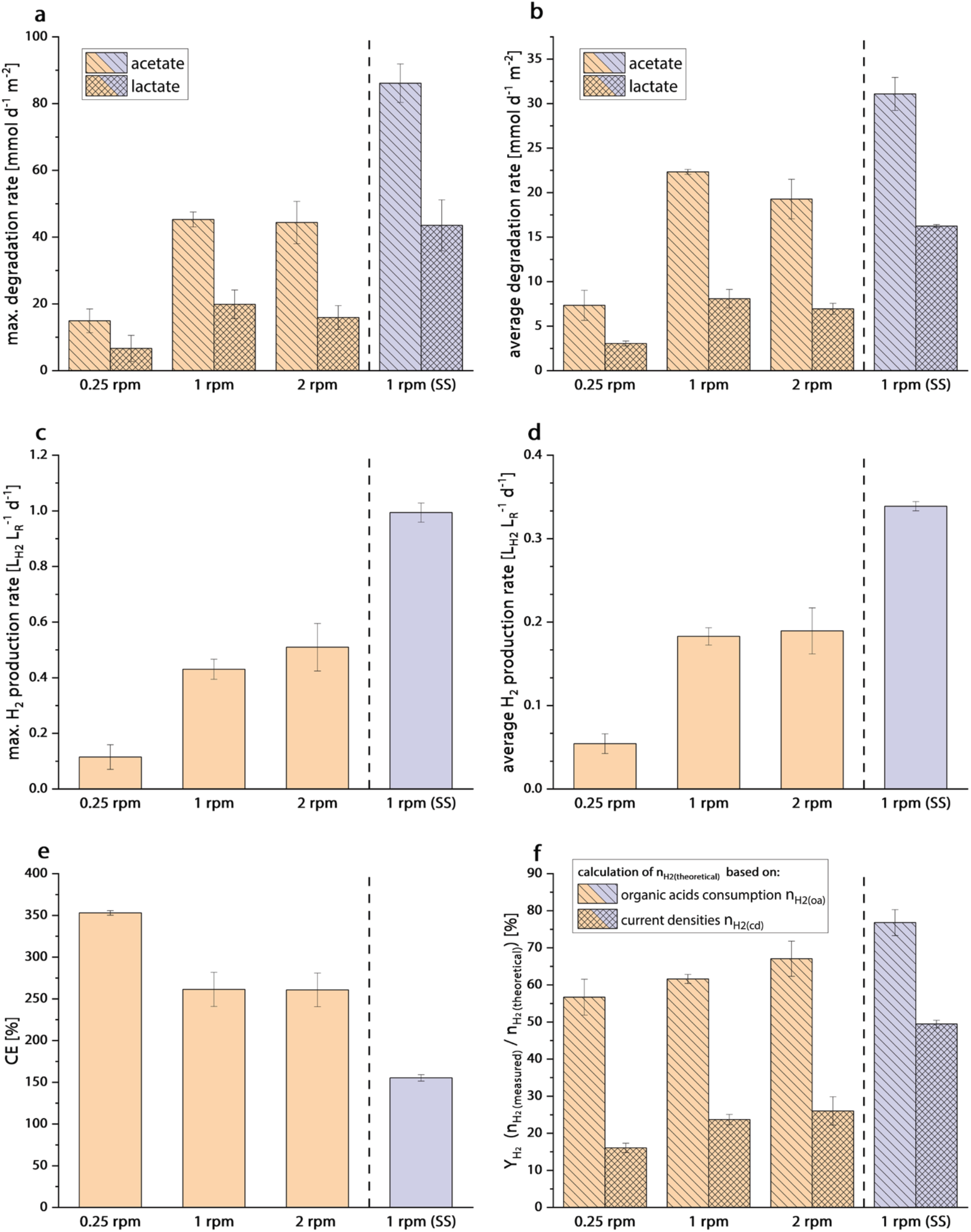
Impact of the rotational speed of the anodes on different performance parameters of the RDBER operated as microbial electrolysis cell (orange). The lilac bar represents the performance of the reactor system operated with the optimized stainless-steel (SS) counter electrode configuration at 1 rpm. The experimental duration of all experiments shown was 9 days. (a) and (b) maximum and average degradation rates of the two organic acids acetate and lactate based on the IC measurements. (c) and (d) maximum and average hydrogen production rates in liters (L_H2_) per liters of reactor volume (L_R_) and day. (e) coulombic “efficiencies” (CE) for the anodic half-cell reaction (ratio of the electrons transferred to the anode and the electrons released from the degraded organic acids). (f) hydrogen yield calculated as the share of the moles of hydrogen measured at the gas outlet of the reactor in the moles of hydrogen that could have been theoretically produced either based on the organic acid consumption (bars filled with lines: n_H2(theoretical)_ = n_H2(oa)_) or calculated from the measured current densities (bars filled with crossed lines: n_H2(theoretical)_ = n_H2(cd)_).

Different approaches are conceivable to reduce or even prevent this phenomenon of hydrogen recycling in a microbial electrolysis cell. The apparently simplest solution is to install a semi-permeable membrane between the anode and cathode compartments. However, this is contrary to one of the main concepts of the RDBER, according to which the high ohmic losses and the complexity of the reactor, especially in the context of future scaling, are to be reduced by the elimination of the membrane. Moreover, biotic hydrogen re-oxidation might also be suppressed by the addition of specific inhibitors. For example, conjugated oligo-electrolytes were added to a membrane-less MEC, which increased hydrogen recovery by 67-fold and acetate removal by 4.4-fold as reported by Liu et al. (Liu et al., 2015). However, the regular addition of high-priced inhibitors to a biotechnological process, specifically a continuous process, would be economically unsound. Furthermore, the RDBER is designed especially for biotechnological pure culture applications due to its autoclavability and the theoretical possibility of clean-in-place. Thus, it would also be feasible to rely on other electroactive strains that are either natively (e.g. *Geobacter metallireducens, Desulfuromonas acetexigens* or *Shewanella oneidensis*) or due to genetic modifications not capable of oxidizing hydrogen. However, the limited number of electroactive microorganisms or the lack of a genetic accessibility often restricts microbial strain options. Additionally, preventing the undesired proliferation of hydrogen oxidizing microorganisms in a mixed-species anodic biofilm presents a challenge. Therefore, we aimed for a constructive approach to minimize hydrogen recycling, as it has already been shown for other bioelectrochemical reactor concepts that a well thought-out arrangement of the electrodes and a well-managed directional flow of the fluid and gas streams in a reactor can significantly reduce electrochemical short circuits between the electrodes (Giddings et al., 2015; Guo et al., 2010; Wang et al., 2023).

### 3.2 Operation of the RDBER with a modified counter electrode

In an attempt to minimize hydrogen shuttling, an alternative counter electrode configuration was developed specifically for MEC operation of the RDBER. For this purpose, the counter electrodes, which act as H_2_-producing cathodes during MEC operation, were moved from the lower half of the reactor to the upper part of the reactor and thus above the rotating drum of the working electrodes. The relocation of the cathodes was intended to prevent the cathode-generated hydrogen bubbles from passing the anodic biofilm on their ascent to the gas outlet of the reactor and thus from being reoxidized by the microbes in the anodic biofilm. As part of this process, we also decided to replace the electrode material (MMO-coated titanium) with a more cost effective perforated and bent stainless steel sheet (a simplified detailed photograph of both electron configurations can be seen in Fig. 1). It should also be mentioned at this point that the basic reactor structure, including the counter-electrode current collection and mounting system, has remained unchanged by these modifications and therefore either the original or the revised counter-electrode arrangement can be installed, depending on the application.

We conducted batch cultivations of *G. sulfurreducens* and *S. oneidensis* in the modified RDBER under identical conditions to the previous batch experiments (e.g. 0 mV vs. SHE, 20 mM lactate, 20 mM acetate in BES-Medium) to allow for direct comparison with these prior cultivations. A rotational speed of 1 rpm was selected for the working electrodes, as the previous rpm-comparison experiments indicated that increasing the speed to 2 rpm provided only marginal performance improvement – if at all –, which did not justify the increased energy consumption and wear on the reactors bearings and rotary shaft seals associated with higher rotational speeds.

As illustrated in Fig. 2, the current density profile recorded in these cultivations exhibits a significantly steeper and more exponential increase, reaching a sharp peak at a twofold higher anodic current density of 1.98 ± 0.11 A m^-2^ (corresponding to a volumetric current density of 198 ± 11 A m^-3^, compared to 100 ± 0.07 A m^-3^ for the previous cathode configuration). Following this peak, the current density declines in a wavelike manner to well below 1 A m^-2^. This performance enhancement is similarly reflected in the specific degradation rates. The maximum acetate degradation rate exceeds 80 mmol d^-1^ m^-2^ anode surface (or approximately 0.5 kg COD d^-1^ m reactor volume), compared to ∼45 mmol d^-1^ m^-2^ in the previous configuration. Likewise, the maximum lactate degradation rate doubled, increasing from 19.9 ± 4.6 to 43.5 ± 7.6 mmol d^-1^ m^-2^. Notably, the increased current densities of the modified reactor also correspond to enhanced hydrogen production rates. For instance, the maximum hydrogen production rate more than doubled, rising from 0.43 ± 0.03 to 0.99 ± 0.03 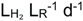. Interestingly, while the average acetate degradation rate increased by approximately 39%, the average hydrogen production rate rose by ca. 90%. This indicates that less hydrogen was reoxidized by the anodic biofilm and that the modification of the cathode, thus, had a positive impact on minimizing the hydrogen shuttling in the system. This effect might also be reflected in the anodic efficiency of the modified system, which is closer to 100%, with a Coulombic efficiency (*CE*) of 155 ± 4% (previously 261 ± 20%). On the one hand, this shows that the contribution of H_2_ as an additional electron donor (that was not considered when calculating the *CE*) could be minimized. On the other hand, it implies that at least one-third of the total electrons transferred to the anode by the anodic biofilm could still originate from hydrogen reoxidation.

According to the calculated 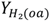, approximately 75% of the electrons theoretically released through organic acid oxidation were recovered in the produced hydrogen. However, this only accounts for about 50% of the expected hydrogen yield based on the measured current densities 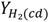. This suggests that while the cathode modification reduced the amount of hydrogen shuttling to some extent, it could not entirely prevent it.

### 3.3 Optical coherence tomography of anodic biofilms

In order to obtain structural information about the anodic biofilm and especially in order to elucidate, whether or not the improvement of the reactor’s performance parameters after the cathode modifications (e.g.: higher current densities) might be a consequence of a change in biofilm quantity or morphology, we conducted an OCT analysis of the biofilm on the foremost anode disc after each batch experiment. Since space-time-resolved in vivo monitoring of biofilm development in the RDBER was already the focus of other studies, and pausing electrode rotation to capture OCT images (which can take up to an hour) could have interfered with the experimental outcome, we conducted only an endpoint analysis of the biofilm in this study - prior to reactor disassembly and cleaning. The biofilm parameters, average biovolume (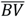 in µm^3^ biofilm per m^2^ anode surface) and substratum coverage (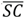 in % of anode surface that is covered with a biofilm) were then calculated from the OCT-data sets (Wagner and Horn, 2017). Fig. 4 suggest that increasing the rotational speed from 0.25 to 1 rpm in the original reactor configuration may enhance biofilm formation, though the large error bars warrant caution in interpretation. However, the bar charts in Fig. 4 clearly show that cathode modification did not significantly increase biovolume or anode coverage, indicating that these factors do not explain the observed rise in current densities or degradation rates due to the reactor modification.

**Figure 4:**
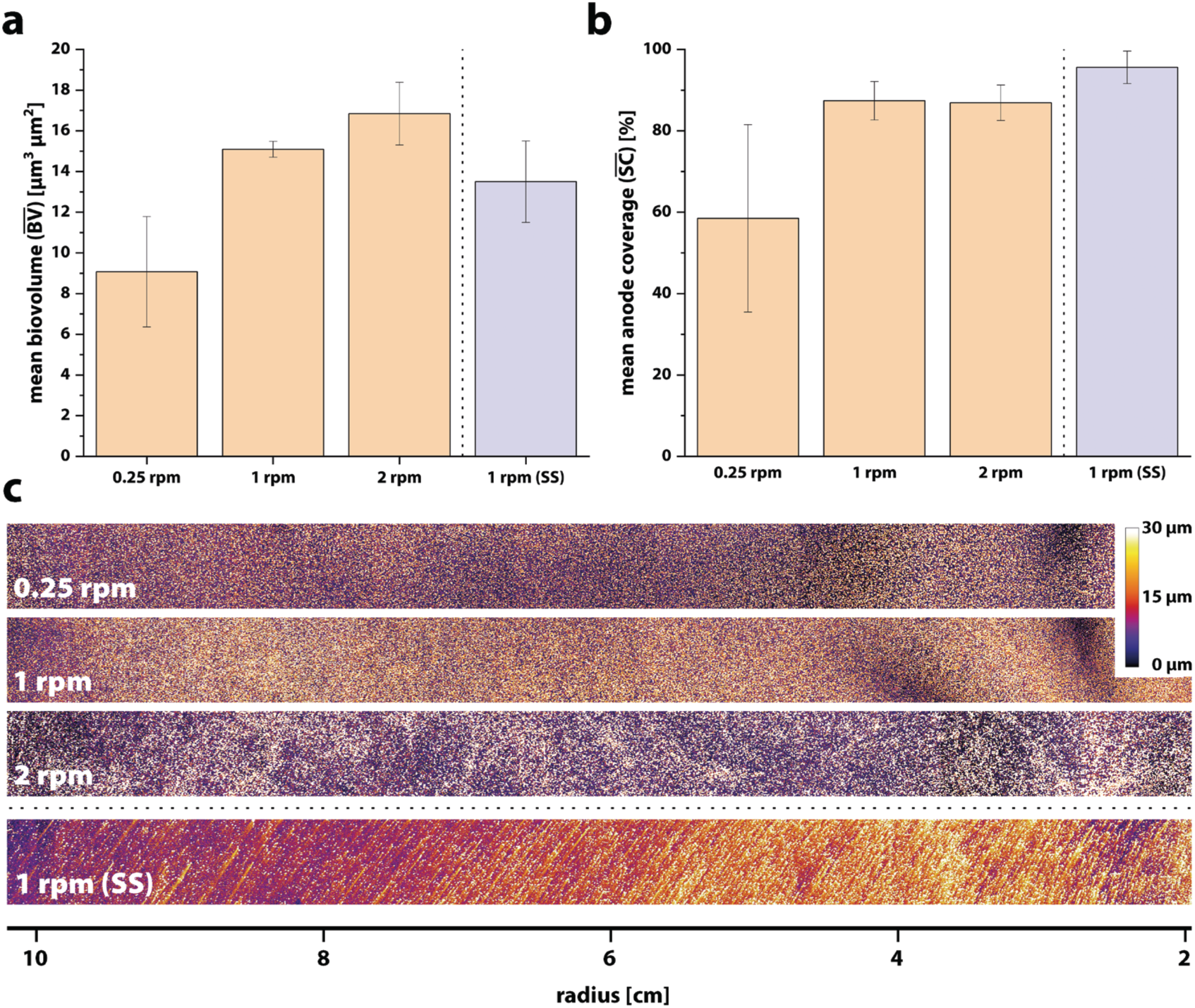
OCT-derived biofilm parameters of the anodic biofilm after 9 days of operation. (a) mean biovolume and (b) mean substratum coverage. The lilac bar represents the biofilm parameters obtained from the experiments conducted with the optimized stainless-steel (SS) counter electrode configuration. (c) OCT-derived height maps of a radial excerpt of the anode after 9 days of operation. The same radial section has been visualized on the working electrode in each case. Colors are coded from black (0 µm) to white (30 μm).

However, it should be mentioned at this point that the interpretation and in particular the comparison of OCT data between different reactor systems must generally be carried out very carefully and should not be over-interpreted. The differences in the counter electrode configuration of the two setups tested in this work may result in different shear forces on the anodic biofilm and thus lead to a different resulting biofilm density, with a higher number of catalytically active cells per biofilm volume. In addition, a denser biofilm could lead to a different diffusional behavior within the biofilm, which could impair the nutrient supply within the biofilm.

However, notably the biofilm morphology undergoes an interesting transformation as a result of the cathode modification. In the original configuration of the RDBER, the anodic biofilms on the foremost graphite disc primarily consist of punctiform, tower-like structures. After the modification, however, distinct shooting star-like formations emerge, with tail angles that vary depending on the disc radius. The underlying causes of this morphological change remain unclear. Potential contributing factors include: (I) reduced disturbance due to fewer passing hydrogen bubbles, (II) a shift in substrate utilization from hydrogen to organic acids, or (III) solely the altered hydrodynamics resulting from the displacement of the counter electrode. Moreover, further experiments are required to determine how homogeneously the biofilm distributes across the 14 different working electrode discs in the reactor and whether growth significantly differs between discs due to suboptimal hydrodynamic conditions, that could lead to hydrodynamically isolated regions and therefor in the most detrimental case, the presence of stagnant dead zones in the reactor. A hydrodynamic model, combined with an in-depth biofilm analysis using molecular biological, optical, and gravimetric methods on all electrode discs during reactor disassembly, will be essential for addressing these questions and further optimizing the system. The overall hydrodynamic design of this version of the RDBER can certainly be improved, as it was not a primary focus during the construction of this first reactor version. For instance, Introducing the recirculated substrate laterally, in addition to the front inlet, and increasing the number of perforations in the graphite discs could improve the mixing of the bulk phase in the gaps between the individual anode discs. This would enhance substrate delivery to these areas, reducing the formation of electrochemically induced pH gradients. Additionally, it would minimize the formation of a ‘laminar-like’ flow around the rotating electrode drum, ensuring better flow through the interelectrode space.

### 3.4 Comparison of the performance of the RDBER operated as a MEC scale-up attempt with other scale-ups

For a comprehensive comparison of the RDBER’s performance with other scale-up attempts, we adopted, modified, and expanded a spreadsheet from a recent literature review by Jiang et al., which systematically analyzed MEC scale-up approaches and their performance (Jiang et al., 2023). As a result, Table 1 compiles construction details and performance parameters from MEC pilot studies with reactor volumes of 10 L or more. To provide a clearer visualization of the performance comparison, we plotted selected performance indicators and reactor characteristics against each other in three different scatter plots (see Fig. 5).

**Figure 5:**
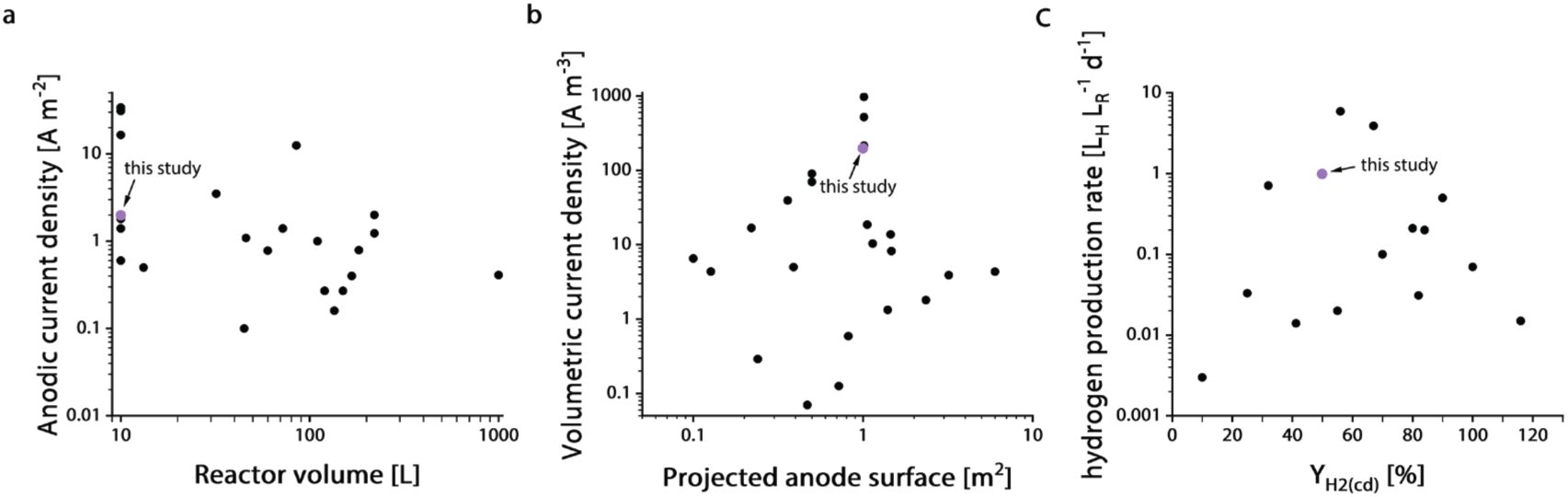
Classification of performance characteristics of the RDBER (lilac dot) in comparison to pilot-scale MEC reactors (≥10 L reactor volume) from other studies (data points derived from *Table 1*). (a) Maximum anodic current densities reported in the literature are plotted against the respective total reactor volume. (b) Maximum volumetric current densities achieved are shown as a function of the projected anode surface. (c) Volumetric hydrogen production rates (L H_2_ per L reactor volume per day) are plotted against the hydrogen yield, calculated from recorded current densities. The hydrogen yield, YH_2_(cd), corresponds to the cathodic efficiency, rcat in percent.

**Table 1:**
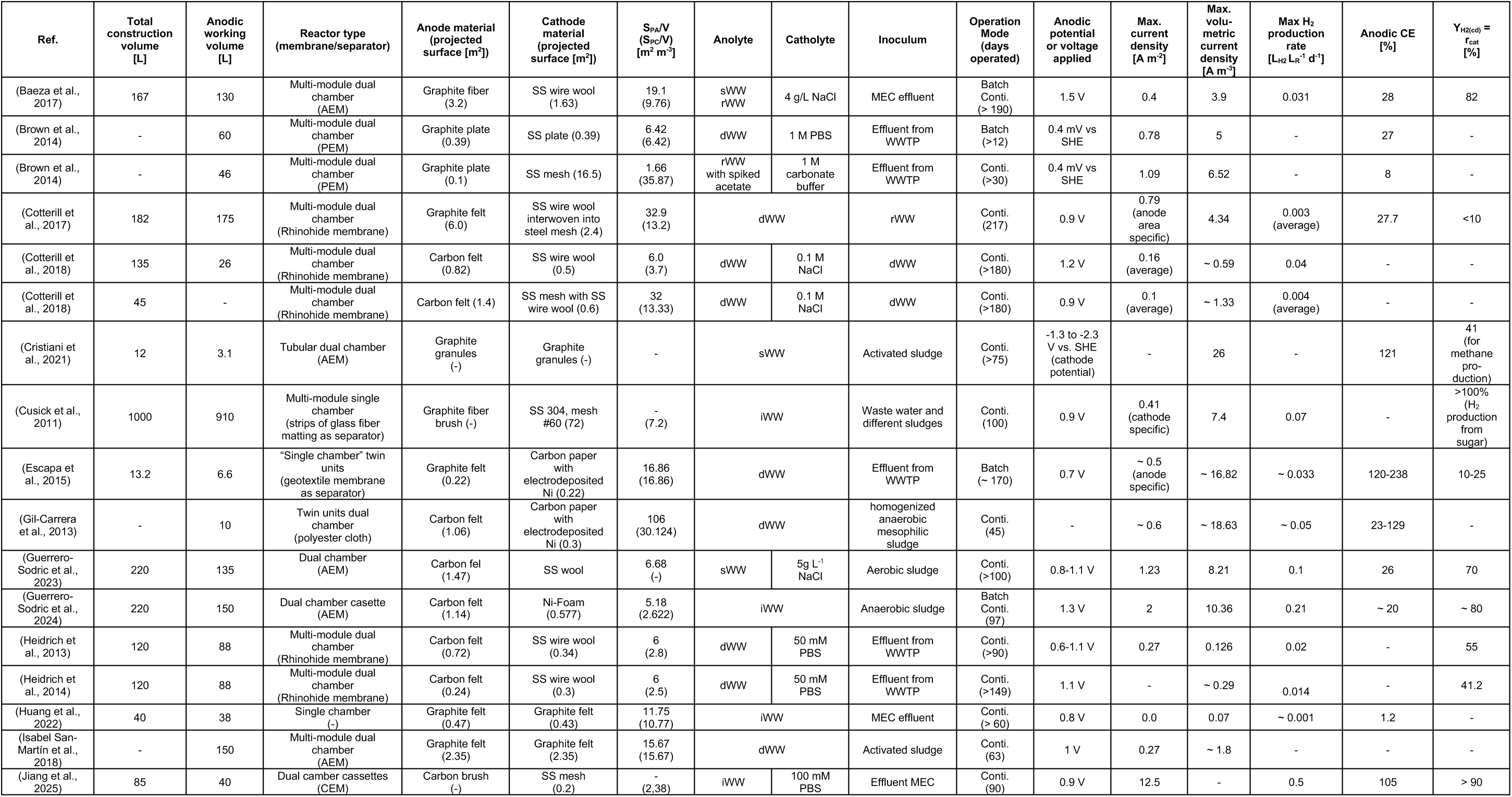
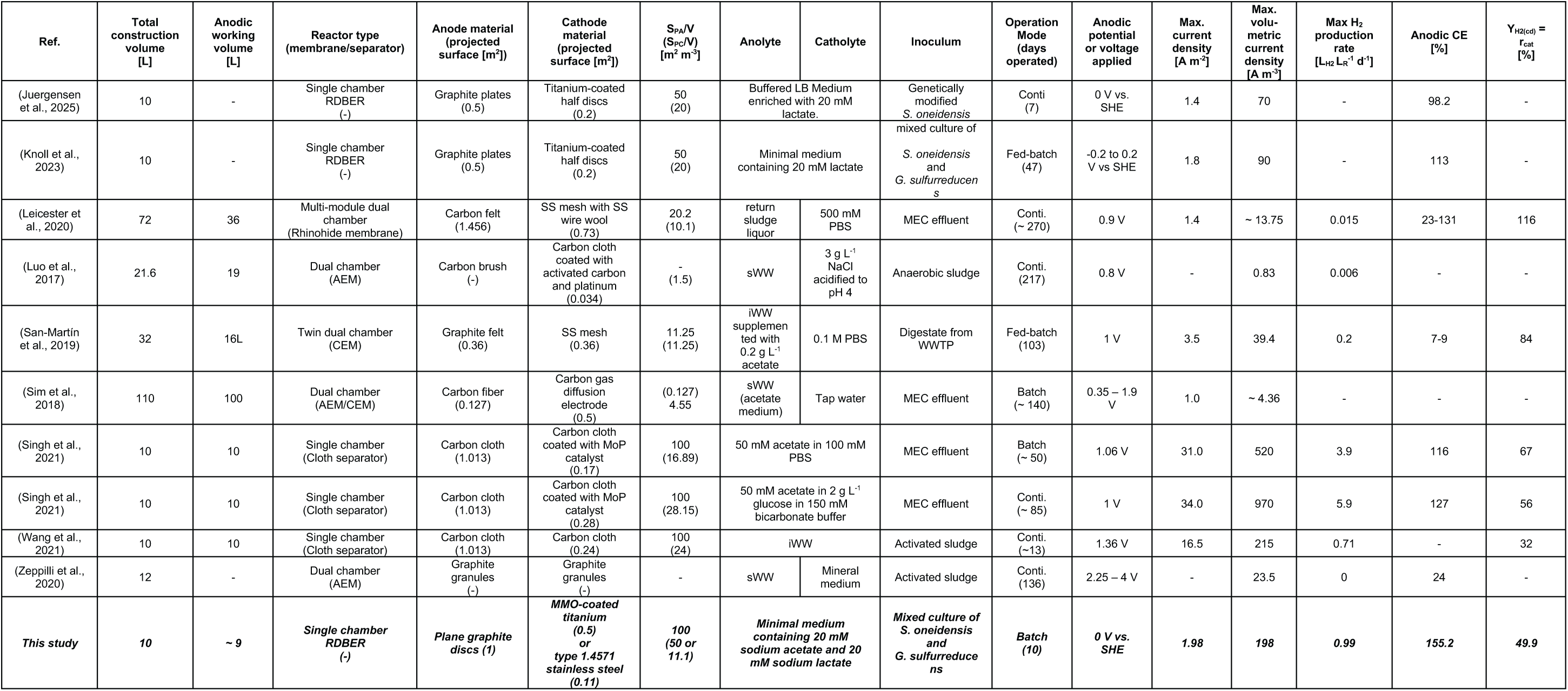
Tabular summary of MEC pilot studies with ≥ 10 L reactor volume, reactor details and performance indicators. AEM – anion exchange membrane, PEM – proton exchange membrane, CEM – cation exchange membrane, S_PA_ – projected anode surface, S_PC_ – projected cathode surface, conti. – continuous, SHE – standard hydrogen electrode, sWW – synthetic waste water, rWW – real waste water, iWW – industrial wastewater, dWW – domestic wastewater, WWTP – waste water treatment plant. List adopted, modified and extended from Jiang et al. 2023(Jiang et al., 2023).

As shown in Table 1, the MEC reactors used in different studies vary not only in obvious parameters such as reactor volume and projected anode or cathode surface area but also in electrode materials and configurations. These differences lead to variations in electrode porosity and surface properties, which in turn influence both electrocatalytic performance and biocompatibility. Moreover, the composition of the anolyte and catholyte differs significantly across studies, ranging from nutrient- and substrate-poor, raw municipal wastewater to industrial waste streams with high organic loads, as well as synthetic, well-buffered media, such as the minimal medium used in this study. Furthermore, the reactors in these studies were operated under different temperature conditions, which can significantly impact MEC performance. Additionally, the mode of operation - whether batch, fed-batch, or continuous - along with feeding parameters such as organic loading rate and hydraulic retention time, can play a crucial role in substrate availability and, consequently, the development of the anodic biofilm.

As a result, such comparisons are inherently difficult - if not nearly impossible - to interpret quantitatively. Nevertheless, they can still offer valuable insights into the strengths and weaknesses of a reactor concept and help guide further optimization efforts. As indicated in Table 1, most published MEC scale-up attempts rely on multiple dual-chamber reactors, often arranged in a cassette-style configuration as part of a numbering-up strategy. Only 8 out of 28 pilot-scale studies with reactor volumes of 10 L or more utilize single-chamber reactors, three of which describe either the reactor examined in this study or a variation thereof. In terms of reactor volume, the 10 L RDBER presented in this study represents the lower end of pilot-scale MECs plotted in Fig. 5a, which includes reactors up to 1000 L. However, scaled up 100 L versions of the 10 L RDBER, each with a 10 m^2^ working electrode surface, are currently being demonstrated and evaluated in different pilot test setups (Pott, 2023).

As shown in Figure 5a, the maximum achieved anodic current density (∼2 A m^-2^) falls within the upper mid-range. In comparison, state-of-the-art flat-plate zero-gap reactors at the laboratory scale (with only a few milliliters of anode working volume) have achieved record current densities of up to 80 A m^-2^ - 40 times higher than the anodic current densities measured in this study. Yet, the RDBER’s current densities are well in range with the anodic current densities obtained by Knoll and Juergensen in a very similar variation of our 10 L RDBER (1.8 A m^-2^ and 1.4 A m^-2^ respectively) (Juergensen et al., 2025; Knoll et al., 2023).

Among the upscaling attempts listed in Table 1, the reactor systems developed in the Liu lab approach for a MEC pilot comparably high current densities of up to 34 A m^-2^. Interestingly, all of these reactors are single-chamber MECs (Singh et al., 2021; Wang et al., 2021). In order to target anodic current densities of similar magnitude in the 10 L RDBER, several approaches are conceivable: As explained in detail in section 3.3, a well-designed and computationally validated flow regime could play a crucial role in improving mixing in the reactor, thereby minimizing dead zones that could otherwise hinder the development of biofilms on distinct areas of some anode discs and thereby limiting the anodic current density.

The plain graphite electrodes currently used in the reactor are an unpopular choice as anode materials in pilot-scale MECs. In the original RDBER design, they were primarily selected for their highly planar surface, which facilitates precise quantitative OCT biofilm analysis. Enhancing the specific surface area of the anode could be achieved by roughening the graphite discs, utilizing alternative porous materials, or integrating graphite felt on a support structure as working electrode discs.

Lastly, reducing the distance between the anode and cathode in the RDBER by replacing the individual rotating graphite discs, which currently serve solely as working electrodes, with zero-gap assemblies separated by functional spacers, could be an interesting approach: These assemblies could be arranged alternately, forming cathode and anode compartments between the discs along the rotating electrode drum. In addition to providing structural stability to the zero-gap assembly discs, the spacers could also guide the bulk flow - driven by their rotation - from the anode “compartments” into the cathode “compartments” (where the medium and gas outlets must be located), thereby preventing hydrogen shuttling.

With a projected anode surface area of 1 m2, the 10 L RDBER falls within the mid-range of pilot-scale MECs in terms of working electrode surface (see Fig. 5b). However, when considering volumetric current densities, the RDBER ranks among the highest-performing systems (ca. 200 A m^-3^). Only the single-chamber reactors from the Liu Lab exceed the RDBER’s performance, though their volumetric current densities remain within the same order of magnitude (215 – 970 A m^-3^) (Singh et al., 2021; Wang et al., 2021). This strong performance is primarily attributed to the RDBER’s high surface-to-volume ratio of 100:1. Consequently, its maximum volumetric hydrogen production rate of approximately 1 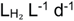 ranks among the top three reported in the literature for MEC pilots of at least 10 L reactor volume, surpassed only by the Liu Lab’s systems, which could achieve 3.9 and 5.9 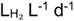.

Finally, Fig 5c shows, that the best hydrogen yield based on current density 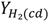 (cathodic hydrogen recovery) reached in this study was approximately 50%, which is lower compared to other reactor systems. This can be explained by the prevalence of dual chamber designs in most of the other studies, which effectively minimize hydrogen shuttling. However, it is worth noting that some of the studies in Table 1 which report exceptionally high 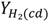 values involved glucose supplementation or formation in the anolyte. Under anoxic conditions, glucose can be metabolized directly in the bulk phase, leading to additional hydrogen production that inflates yield calculations. Interestingly, even the RDBER’s average hydrogen production rate of 0.5 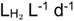 remains higher than most maximum hydrogen production rates reported for pilot scale MECs studies listed in Table 1.

## 4. Conclusions

Without significant advancements in optimizing the reactor’s key performance parameters, the RDBER - despite currently outperforming most other MEC scale-up attempts - may eventually be outpaced by the continued development of zero-gap-based systems employing a numbering-up strategy. However, while the scalability of such systems is frequently framed as a mere engineering challenge it remains to be demonstrated that stackable modules can effectively process hundreds or even thousands of liters of wastewater streams. For applications requiring a large anodic working volume, whether due to high wastewater throughput or a substantial presence of suspended solids, a system like the RDBER could still offer unique advantages.

## CRediT authorship contribution statement

**Zhizhao Xiao:** Investigation, Writing - Review & Editing **Max Rümenapf:** Investigation, Writing - Review & Editing **Max Hackbarth:** Investigation, Conceptualization, Methodology **Andrea Hille-Reichel:** Conceptualization, Writing - Review & Editing **Harald Horn:** Conceptualization, Writing - Review & Editing, Funding acquisition. **Johannes Eberhard Reiner:** Investigation, Conceptualization, Methodology, Visualization, Writing - Original Draft & Editing.

## Data availability

Data will be made available on request.

## Acknowledgements

Special thanks to Axel Heidt as the person responsible for the IC measurements and to Erwin Wachter, Joachim Lang and Alfred Herbst from the metal workshop of the EBI for their efforts and support.

