## Supplementary Information for "Impact of the rotational speed and counter electrode configuration on the performance of a rotating disc bioelectrochemical reactor (RDBER) operated as microbial electrolysis cell"

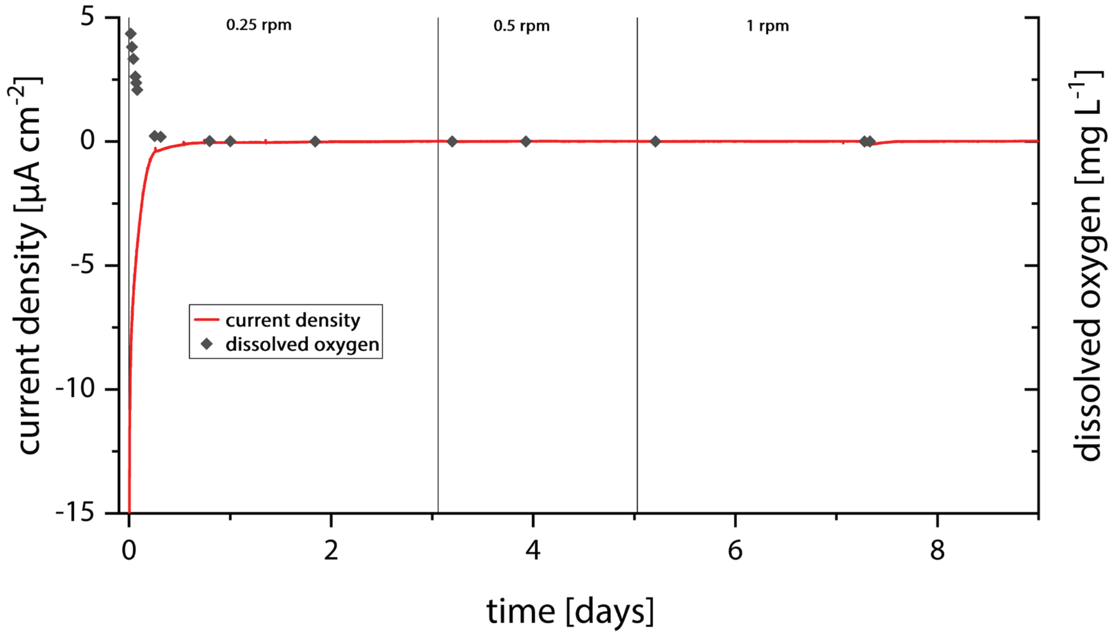


*Supplementary Figure 1:* Current density and dissolved oxygen in an abiotic control experiment. No degradation of the organic acids could be detected in the ion chromatography data.
